# Spatiotemporal characterization of the neural correlates of outcome valence and surprise during reward learning in humans

**DOI:** 10.1101/091710

**Authors:** Elsa Fouragnan, Filippo Queirazza, Chris Retzler, Karen J. Mullinger, Marios G. Philiastides

## Abstract

Reward learning depends on accurate reward associations with potential choices. Two separate outcome dimensions, namely the valence (positive or negative) and surprise (the absolute degree of deviation from expectations) of an outcome are thought to subserve adaptive decision-making and learning, however their neural correlates and relative contribution to learning remain debated. Here, we coupled single-trial analyses of electroencephalography with simultaneously acquired fMRI, while participants performed a probabilistic reversal-learning task, to offer evidence of temporally overlapping but largely distinct spatial representations of outcome valence and surprise in the human brain. Electrophysiological variability in outcome valence correlated with activity in regions of the human reward network promoting approach or avoidance learning. Variability in outcome surprise correlated primarily with activity in regions of the human attentional network controlling the speed of learning. Crucially, despite the largely separate spatial extend of these representations we also found a linear superposition of the two outcome dimensions in a smaller network encompassing visuo-mnemonic and reward areas. This spatiotemporal overlap was uniquely exposed by our EEG-informed fMRI approach. Activity in this network was further predictive of stimulus value updating indicating a comparable contribution of both signals to reward learning.

## Introduction

Learning occurs with the combination of two critical dimensions of decision outcome signaling discrepancies between expectations and reality: the categorical valence (i.e. whether an outcome is overall better or worse than expected [positive or negative]) and surprise, or the degree of deviation from expectations ([high or low]). Traditional learning theories ^1,2^ have treated these quantities as two facets of a single unified representation, commonly referred to as the reward prediction error (RPE) signal. Correspondingly, many animal electrophysiology ^3,4^ and human neuroimaging studies ^5,6^ have focused on identifying brain areas exhibiting a positive relationship with a fully parametric (signed) RPE signal. To date, however, the extent to which outcome valence and surprise (unsigned RPE) ^7,8^ can also be encoded separately in the human brain and the degree to which they converge (in space and time) to drive learning, remain poorly understood.

Investigating these open questions is critical as outcome valence and surprise could subserve related but largely separate functions. For instance, the valence of an outcome determines the direction in which learning will occur. Positive outcomes will increase the likelihood of similar decisions in the future, whereas negative outcomes will increase the likelihood of an avoidance response ^9-11^. In contrast, the surprise dimension modulates the amount of attention devoted to outcomes, which in turn dictates the degree of learning and ultimately the updating of future reward expectations. Highly unpredicted outcomes (either positive or negative) can boost attention and facilitate learning, whereas less surprising outcomes attracting reduced attention might slow down the learning process instead ^12,13^.

In line with these separate functional roles, recent studies in animals and humans have observed distinct neural signals aligning both with outcome valence and surprise within a distributed network of areas, including the midbrain ^l4,15^, the striatum ^16-18^, the cingulate cortex and the anterior insula ^7,19-21^. Specifically, these recent studies employing model-based fMRI have begun to characterize the neural basis of both signals providing new evidence that separate brain networks might encode outcome valence and surprise in the human brain ^22,23^. Given the poor temporal resolution of fMRI, however, these studies preclude a rigorous assessment of the relative timing and underlying dynamics of these two learning-related signals.

Our previous electroencephalography (EEG) work revealed two distinct components encoding categorical outcome valence (early and late) and a separate outcome surprise component that overlapped temporally with the later of the two valence components ^24^, suggesting that the brain might require simultaneous access to both signals to drive learning. The poor spatial resolution of the EEG, however, prevented a thorough characterization of the spatial generators associated with each outcome representation. More recently, we used simultaneous EEG-fMRI, to map out the spatiotemporal dynamics of the two outcome valence components ^11^. However, no analysis was presented that examined the neural systems associated with surprise or their relationship with outcome valence.

Here, using the data in ^11^ we aim to provide a comprehensive spatiotemporal characterization of the relationship between the late temporally overlapping representations of outcome value and salience. Our hypothesis is that endogenous trial-to-trial variability in electrophysiologically derived measures of the two outcome variables can be used to form separate fMRI predictors to tease apart the brain networks associated with each representation. Moreover, we directly test the hypothesis that temporal and spatial congruency, of otherwise separate outcome value and salience representations, constitutes a viable mechanism for driving reward learning.

## Results

To test our hypotheses and investigate the relative and potentially combined contribution (spatial and temporal) of outcome valence and surprise to learning, we present new analyses of data reported in ^11^. Twenty participants were engaged in a probabilistic reversal-learning task ^25^ while we recorded simultaneous EEG-fMRI data. Specifically, on each trial subjects were presented with two abstract symbols and through feedback learned to select the one with the highest reward probability. After reaching a predefined learning criterion, the high reward probability was re-assigned to a different symbol and subjects had to enter a new learning phase (see Fig. 1a and Materials and Methods). Participants’ behaviors were probabilistic and mirrored the principles of a RL mechanism ^1^ (Fig. 1b) best explained with a simple model-free RL with a fixed learning rate (see model comparisons in Materials and Methods).

Temporal superposition of outcome valence and surprise. To identify temporal components encoding outcome valence and surprise we used multivariate discriminant analysis of feedback-locked EEG signals ^26-28^. Specifically, for each participant, we estimated linear weighting of the EEG electrode signals that maximally discriminated between conditions of interests (see below) over several distinct temporal windows (Eq. 5). Applying the estimated electrode weights to single-trial data produced a measurement of the discriminating component amplitudes (henceforth *y*), which represent the distance of individual trials from the discriminating hyperplane (Fig. 2a). We later use the single-trial variability (STV) in these values to construct parametric fMRI regressors to identify the brain regions that correlate with the resultant discriminating EEG components (Fig. 2b). To quantify the discriminator’s performance over time we used the area under a receiver operating characteristic curve (i.e. A_z_ value) with a leave-one-out trial cross validation approach.

**Figure 2:**
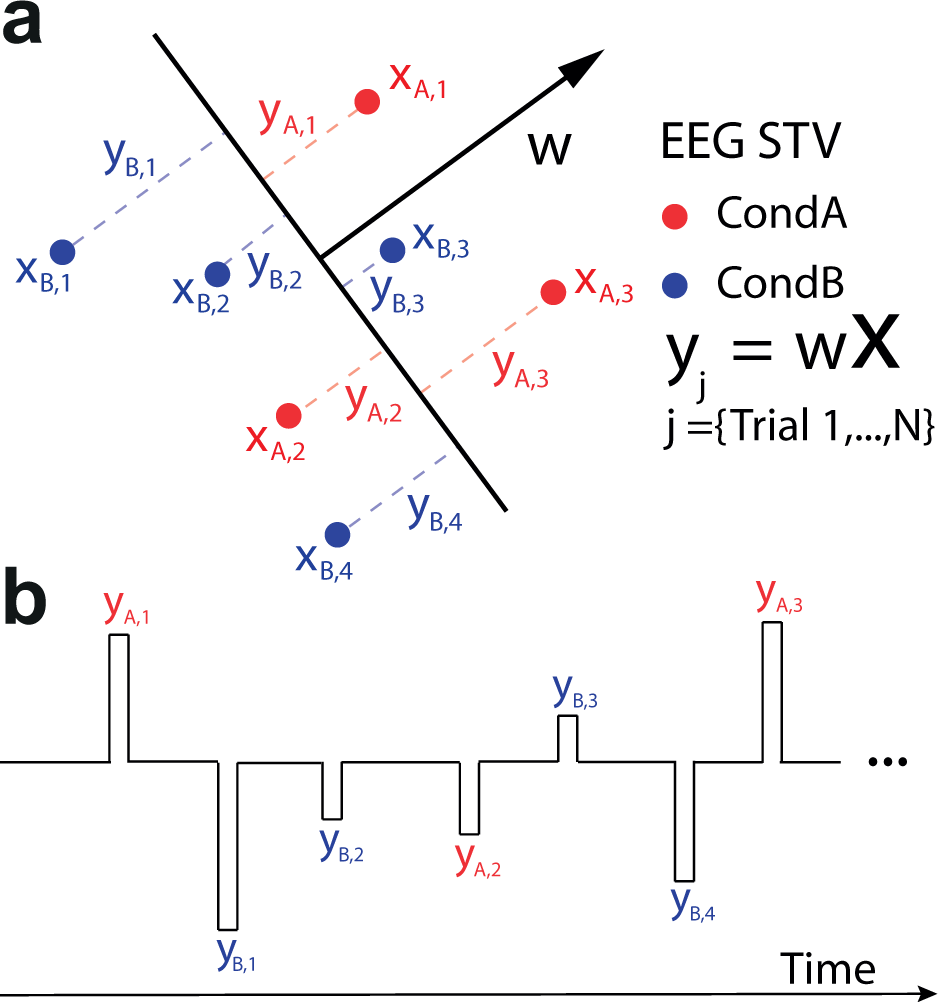
Single-trial discriminant analysis and EEG-informed fMRI regressors. (**a**) We used single-trial analysis of the EEG to perform binary discriminations between conditions of interest, here denoted as condition A and B (in red and blue respectively). We first estimated w, which is a linear weighting on the EEG sensor data (*X*) that maximally discriminates between the two conditions. This determines a task-related projection (*y*) of the data, in which the distance to the decision boundary reflects the decision certainty of the classifier in separating each of the relevant conditions. We treated the single-trial *y* amplitudes (single-trial variability [EEG STV]), as an index of how each condition of interest was perceived on individual trials. (**b**) Given these y values and their corresponding outcome-locked onset time points, we built fMRI regressors for subsequent GLM analyses. These regressors were all convolved with the canonical HRF. Details of specific events included in each EEG-informed fMRI regressor can be found in the main text (see *fMRI analysis* section).

Using this approach, we previously established the presence of two temporally specific EEG components discriminating – reliably in individual participants – between positive and negative outcomes in the time range 180-370 ms after an outcome (Fig. 3a) ^11^. The second of those valence components peaked on average 308 ms (SD ± 37.7) after outcome and was the only component that was directly linked to reward learning by either up- or down- regulating the human reward network to update the expected value of the stimuli following positive and negative outcomes respectively (the first valence component reflected an early alertness response and was selective for negative outcomes only). Importantly, both of these outcome valence components were decoupled (i.e. uncorrelated) from outcome surprise as reported in ^11^.

In this work, we ran a separate multivariate analysis of the EEG to directly discriminate along a outcome surprise dimension, by discriminating between trials with very high (in the range [0.8 – 1]) and very low (in the range [0 – 0.2]) unsigned RPE. In doing so, we identified a separate EEG component that peaked, on average, at 320 ms (SD ± 32.1) after outcome (Fig. 3b). The timing of this component overlapped with that of the second outcome valence component presented above (see shaded regions in Fig. 3a and 3b). We formally tested if the peak times of the two EEG components were significantly different across individuals and found no significant differences (paired t-test, t_19_ = 1.05, P = 0.31), suggesting that the brain might require near simultaneous access to both signals to drive learning.

To formally test whether this additional EEG component was parametrically modulated by outcome surprise (rather than responding categorically to high vs. low surprise – unsigned RPE), we computed discriminator amplitudes for trials with intermediate surprise levels (i.e. high unsigned RPE [0.6 – 0.8]; medium unsigned RPE [0.4 – 0.6] and low unsigned RPE [0.2 – 0.4]), which were not originally used to train the classifier (“unseen” data). Specifically, we applied the spatial weights of the window that resulted in the highest discrimination performance for the extreme outcome surprise levels to the EEG data with intermediate values.

We expected that these “unseen” trials would show a parametric response profile as a function of outcome surprise and therefore the resulting mean component amplitude at the time of peak discrimination would proceed from very low < low < medium < high < very high surprise. Using this approach, we demonstrated that the mean discriminator output (*y*) for each categories parametrically increased as a function of the unsigned RPE signal (ANOVA including all 5 bins: *F*_4,76_ = 43.3, P < 0.001, ANOVA including only the 3 intermediate bins of “unseen” trials: *F*^2,38^ = 14.14, P < 0.001, post hoc paired t-tests carried out for all pairs of consecutive values; all *P* values < 0.005; Fig. 3c, Grey: Intermediate categories, Yellow: Categories used for discrimination), thereby establishing a concrete link between surprise and our EEG component.

Next, we investigated the extent to which the STV in this EEG component predicted the trial-by-trial fluctuations in outcome surprises estimated from our RL model across participants. Importantly, both the outcome surprise estimates and the EEG STV were scaled similarly across participants such that the model estimates were always in the range [0 – 1] and the EEG amplitudes were “normalized” by the discrimination procedure (i.e. centered at zero for all participants). As expected from the analysis above, we found a significant but only moderate correlation between the EEG STV and the single-trial model-based unsigned RPEs (*r* = 0.31, P < 0.001, Fig. 3d). This moderate correlation indicates that the trial-by-trial variability in the EEG amplitudes could offer additional explanatory power in our subsequent fMRI analysis (i.e. in addition to the model-derived estimates of unsigned RPE) to enable a more comprehensive characterization of the brain networks involved in encoding outcome surprise.

To reinforce the notion that the EEG STV in this component carried task-relevant information, which we could exploit further in the fMRI analysis, we performed an additional single-trial regression analysis. Specifically, we showed that the trial-by-trial fluctuations in our outcome surprise component were predictive of the single-trial learning rates in a version of our RL model with a dynamically updating learning rate parameter ^25,29-31^. In other words, the higher the surprise (as indexed by the EEG), the more recent observations were considered for updating the stimulus value for the next decision, thereby boosting the speed of learning and thus increase the single-trial learning rates (*t*_19_ = 3.14, *P* = 0.005). Ultimately, this result offers a more concrete link between behavior and neuronal variability in our EEG outcome surprise component.

**Figure 3:**
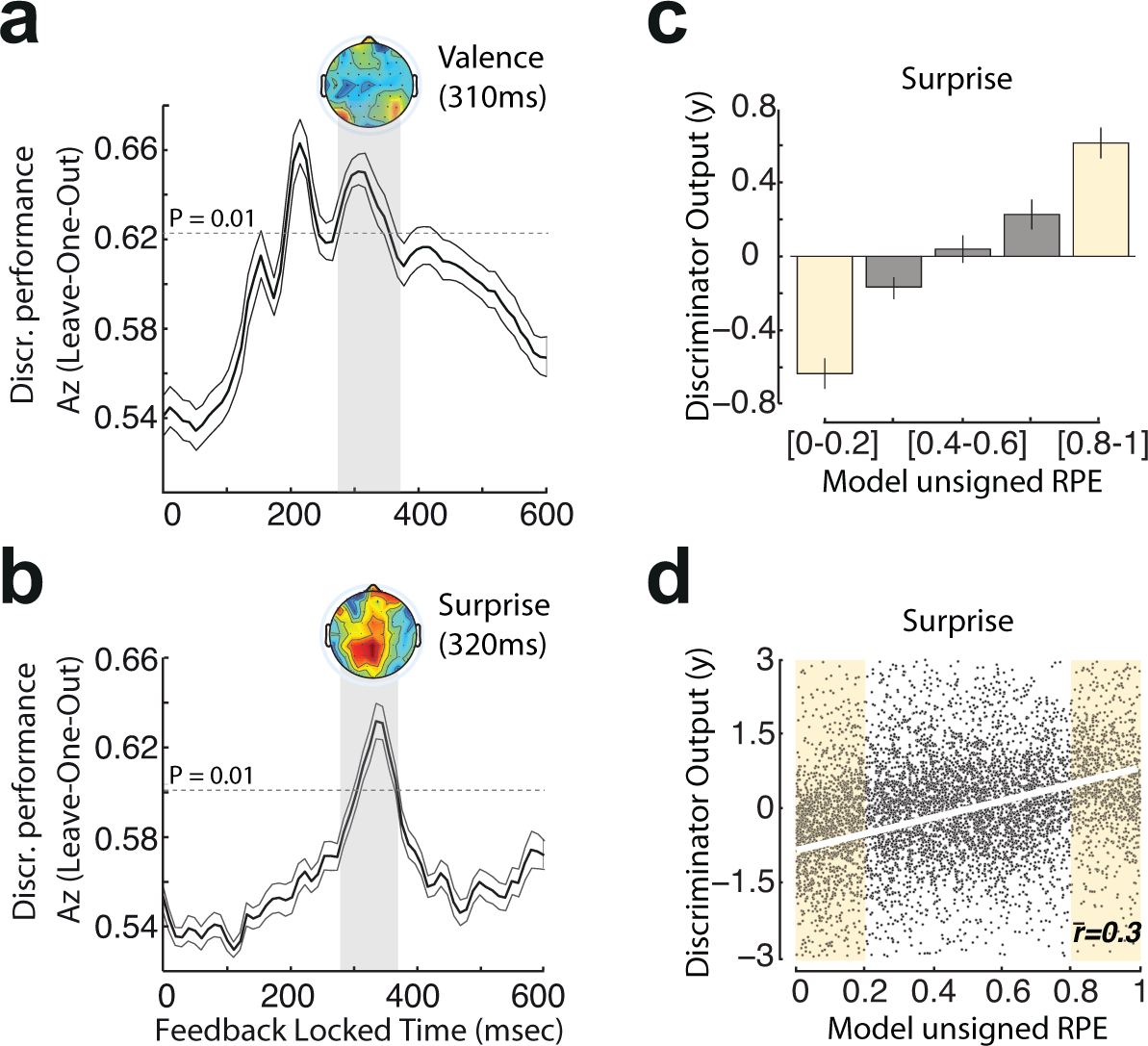
Single-trial EEG analyses. (**a**) Discriminator performance (cross-validated *A*_*Z*_) during valence discrimination (positive vs. negative outcomes) of outcome-locked EEG responses, averaged across subjects (*N* = 20). The dotted line represents the average Az value leading to a significance level of *P* = 0.01, estimated using a bootstrap test. Shaded error bars are standard errors across subjects. In this work, we are focusing only on the second of the two outcome valence components. The scalp map represents the spatial topography of this component. (**b**) Discriminator performance (cross-validated *A*_*Z*_) during outcome surprise discrimination (very low vs. very high surprising outcome), averaged across subjects (*N* = 20). The dotted line represents the average *A*_*Z*_ value leading to a significance level of *P* = 0.01, estimated using a bootstrap test. Shaded error bars are standard errors across subjects. The scalp map represents the spatial topography of the outcome surprise component. (**c**) Mean discriminator output (*y*) for the outcome surprise component, binned in five quantiles based on model-based unsigned RPE estimates, showing a parametric response along the outcome surprise dimension. Yellow bins indicate trials used to train the classifier, while grey bins contain “unseen&” data with intermediate outcome surprises. Error bars are standard errors across subjects. (**d**) Individual trial-by-trial correlations between EEG component amplitudes (*y*) and model-based estimates of outcome surprise (trials used for discrimination are shaded in yellow). Unsigned RPE estimates and the EEG STV were scaled similarly across participants (see Methods). The r bar represents mean correlation coefficient across participants and the white line corresponds to the linear fit through the data.

### Spatial signatures of outcome valence and surprise

Our main fMRI analysis was designed to expose the neural correlaes of the temporally overlapping representations of outcome valence and surprise in a whole-brain analysis by using the explanatory power of the STV in the EEG components associated with these representations. We hypothesize that endogenous variability in these components in individual participants can carry additional information about the internal processing of outcome valence and surprise usually estimated with a simple categorical contrast (positive-vs-negative outcomes) and a model-based unsigned RPE regressor, respectively.

Despite the temporal superposition of the outcome valence and surprise components, their EEG scalp topographies looked qualitatively different (Fig. 3a and 3b), indicating that the two signals might be encoded together in time but in largely different networks. In addition, trial-by-trial amplitude variations in the two discriminating components were largely uncorrelated (*r* = 0.04; *t*_19_ = 0.35; *P* = 0.73), allowing us to use the endogenous STV in the component amplitudes to build parametric, EEG-informed, fMRI regressors to identify the brain networks correlating with each component. Thus, our main GLM included parametric regressors for outcome valence using the STV in the relevant EEG component and parametric regressors for outcome surprise using the model-based unsigned RPE estimates and the STV in the relevant EEG component (see Materials and Methods for a full description of the GLM).

Our EEG-informed fMRI analysis revealed largely separate and distributed networks encoding outcome valence and surprise information. As we demonstrated recently ^11^, the endogenous variability in the valence component covaried with a number of areas of the human reward network ^2,32,33^, such as the ventromedial prefrontal cortex (vmPFC) (Left: 10, −6, 52), the striatum (STR) (Right: 8, 10, −6; Left: −8, 10, −8), the amygdala (Right: 26, −2, −22; Left: −20, −4, −18), the dorsal posterior cingulate cortex (PCC) (Left: −8, −38, 32), the ventral PCC (Left: −2, −50, 20), the putamen (Right: 30, −6, 2; Left: −28, −6, 2), the lateral orbitofrontal cortex (lOFC) (Right: 34, 18, −18; Left: −24, 12, −20) and the posterior insula (INS) (Left: −40, −2, 8) (Fig. 4a for whole-brain results). In addition to these known reward structures, we also identified significant clusters in the lingual gyrus (LG) (Left: −8, −56, 8), the middle temporal gyrus (MTG) (Right: 56, −46, −8; Left: −64, −48, −6) and the precuneus (Left: −10, −56, 30), areas that have previously been implicated in memory retrieval and adaptive visual learning processes ^2^. Activity in this rather distributed network was overall found to be both suppressed and activated in response to negative and positive outcomes respectively, an activity pattern consistent with a role in motivating both avoidance and approach learning^11,34^.

To decipher how the encoding of outcome surprise is represented in the brain relative to the encoding of outcome valence, we used separate parametric fMRI predictors for surprise that were derived from a RL model (i.e. estimated purely from behavior) and EEG STV (i.e. endogenous variability) in our surprise component, respectively. Our conventional model-based fMRI regressor correlated significantly with activity in the dorsolateral PFC (dlPFC) (Right: 48, 12, 34; Left: −50, 8, 38), the bilateral anterior INS (aINS) (Right: 34, 18, 0; Left: −36, 20, 0), the medial PFC (mPFC) (Right: 2, 20, 52), the inferior frontal gyrus (Right: 52, 10, 18), the supramarginal gyrus (Right: 40, −38, −38; Left: −40, −48, 42), the precentral gyrus (Right: 38, 4, 34; Left: −52, 0, 34) and the angular gyrus (Right: 40, −48, 40; Left: −46, −54, 42) (Fig. 4b; yellow clusters), consistent with previous reports ^23^. In particular, activity in the middle frontal gyrus and insular cortex have been consistently linked to deviations from expectations in a large range of learning tasks ^35,36^.

Crucially, using our EEG-informed surprise regressor revealed additional activations over and above what was already conferred by its model-based counterpart (paired t-tests, all *P* < 0.05). These activations included the dorsal STR (Right: 12, 16, 6; Left: −10, 12, 6), the vmPFC (Left: −6, 54, 0), the lOFC (Right: 28, 30, −14; Left: −34, 30, 14) the LG (Right: 12, −62, 12; Left: −10, −60, 8), the bilateral MTG (Right: 60, −32, 2; Left: −64, −30, 2), the occipital pole (Right: 6, −86, 12; Left: −24, −90, 16) and the precuneus (Left: −38, −44, 42) (Fig. 4b; green clusters). As our two regressors for surprise were partially correlated (Fig. 3d) we also repeated this analysis by orthogonalizing our EEG STV regressor with respect to the model-based one (such that any common variance was absorbed by the latter) and obtained similar results. We view this as further evidence that the EEG STV can be used to reveal relevant brain regions and that the approach can be used to complement model-based fMRI.

**Figure 4:**
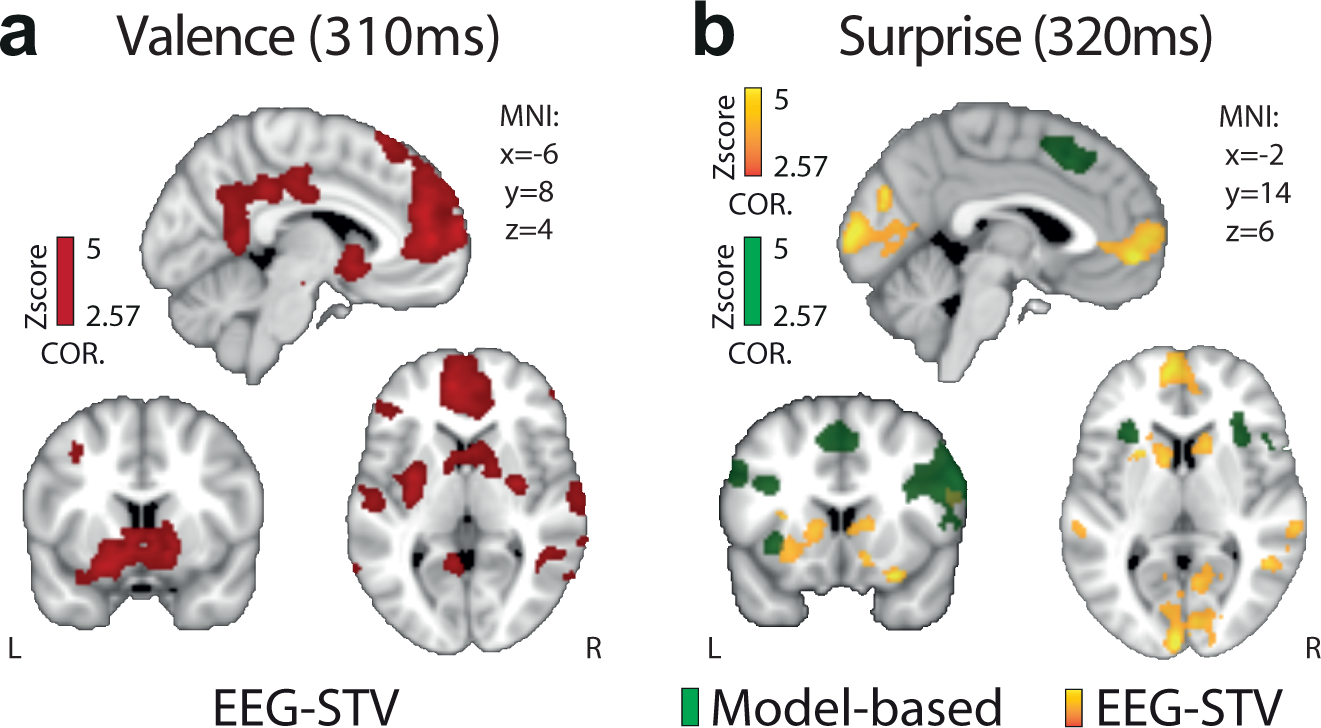
Spatiotemporal characterization of outcome valence and surprise signals. (**a**) Regions correlating with the EEG STV in our valence component, exhibiting overall greater response for positive compared to negative outcomes. (**b**) Regions correlating positively with outcome surprise as captured by a RL model (green) and the STV in our corresponding EEG component (yellow), respectively. Note the complementary nature of activations in the EEG STV map. All activations represent mixed-effects and are rendered on the standard MNI brain at Z > 2.57, corrected using a resampling procedure (minimum cluster size = 76 voxels).

### Temporal and spatial congruency between outcome valence and surprise

Our temporally overlapping representations for outcome valence and surprise suggest that the brain might require near simultaneous access to both signals to drive learning. This notion is also in line with classical RL theory which posits that reward learning is driven by a combination of both categorical valence and unsigned RPE information^3^. We thus, tested whether, within the largely separate spatial representations of valence and surprise (Fig. 4a and 4b), there was a subset of regions in which both outcome dimensions were linearly combined.

To this end, we ran a conjunction analysis across statistical maps identified in our previous EEG-informed GLM. Specifically, among the unique areas found in the outcome surprise network (covarying with the EEG STV), four regions showed a significant overlap with the valence network, namely vmPFC, STR, MTG and LG (see Fig. 5a for the full spatial representation of both networks). While converging evidence highlights the role of the STR and the vmPFC in reward learning ^30^, this result provides new evidence that additional areas such as the MTG and LG are implicated in processing reward learning. Noteworthy is that all four of these regions appeared in the conjunction analysis with our EEG-informed rather than the model-based predictor of unsigned RPE, further highlighting the utility of our simultaneous EEG-fMRI measurements in revealing latent brain states and complementing traditional model-based fMRI approaches. To further validate this point, we ran a separate GLM including only the model-based unsigned RPE and a conventional outcome valence regressor (+1 and −1 for positive and negative outcomes respectively) and conducted a similar conjunction analysis. We found that none of the four regions we identified above appeared in the conjunction analysis of the conventional outcome valence and unsigned RPE regressors (all *P* values > 0.5).

To visualize the signal pattern in these areas, we carried a separate ROI analysis and binned the trials in six categories based on outcome valence (positive or negative outcomes) and surprise (low, medium or high based on our EEG STV of the corresponding component). Note that this analysis was performed for illustrative purposes only (the ROIs were formally linked to outcome valence and surprise in the previous EEG-informed fMRI analysis). We found that activity in these regions exhibited a profile consistent with a linear superposition of the two outcome dimensions with the fMRI signal being overall higher for positive vs. negative outcomes (see illustration in Fig. 5a), while within each of the two outcome types was further modulated by surprise (Fig. 5b). This linear superposition is noteworthy because it points to a common network for the expression of both outcome valence and surprise. This interpretation is consistent with previous evidence that separate families of intermixed neurons are expressed within the same brain regions in response to both outcome valence and unsigned RPE^14^.

To offer additional evidence linking these areas with reward learning, we performed an additional analysis. Specifically, we hypothesized that activity in a network reflecting both outcome valence and surprise should drive learning by encoding the value update required for the chosen option. To test for this formally we used the percent signal change (PSC) in each of the four brain areas as a predictor of expected value updates estimated with our RL model in a linear regression model (Eq. 9). This analysis revealed that all regions were significantly predictive of the update in expected value at time of outcome (vmPFC: *P* < 0.01, *t*_19_ = 2.79, STR: *P* < 0.05, *t*_19_ = 2.06, LG: *P* < 0.001, *t*_19_ = 3.78, MTG: *P* < 0.03, *t*_19_ = 2.29).

**Figure 5:**
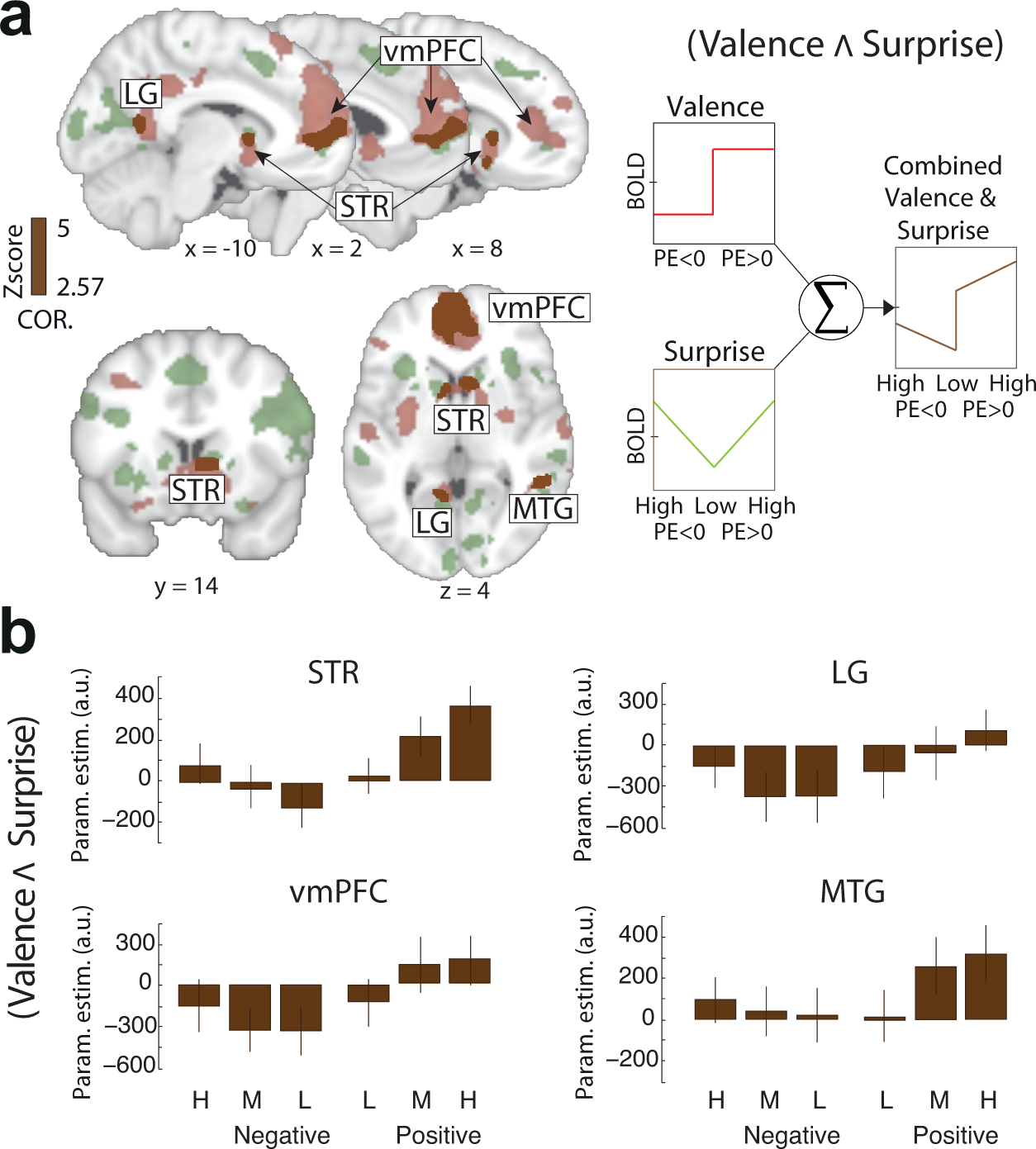
Full spatial representation of the outcome valence and surprise networks and their overlap. (**a**) A conjunction analysis on the results arising from the EEG-informed regressors for the outcome valence and surprise components revealed that four areas – the STR, vmPFC, LG and MTG – significantly encoded both quantities. The conjunction analysis was performed using the resulting whole brain activation maps for outcome valence and surprise and applying a Z > 2.57 and cluster corrected at *P* < 0.05. Results are shown in brown. (**b**) The four overlapping regions exhibited a clear superposition profile between the two signals with a higher BOLD signal for positive vs. negative outcomes but also a systematic increase from low (L), to medium (M), to high (H) outcome surprise trials, within each outcome type.

### Network dynamics underlying outcome valence and surprise signals

Having established that outcome valence and surprise signals were converging later in the outcome period on vmPFC, STR, MTG and LG, we performed an exploratory analysis of the dynamics of this network using DCM. We designed several neural models representing anatomically realistic connections between the four areas and considered both feedforward and feedback information propagations between the nodes, when anatomically possible (see Materials and Methods). We thus structured our model space in four model families to determine the best proposition regarding the connectivity between the vmPFC, the STR, MTG and LG. Specifically, each model family was comprised of equal numbers of models (64) which represented different connectivity patterns between vmPFC and MTG (family [i] represented all models with feedforward information propagation from vmPFC to MTG such that vmPFC – > MTG, family [ii] represented all models with feedback propagation between vmPFC and MTG such that MTG – > vmPFC, family [iii] represented all models in which the vmPFC had a bilateral connection with MTG (vmPFC <-> MTG) and family [iv] represented all models in which the vmPFC had no connection with MTG).

On the basis of previous neurophysiological reports and given our assumptions that learning is ultimately triggered by both outcome valence and surprise, the driving inputs representing these quantities (e.g. the EEG STV in the relevant components) were targeting the STR, which is known to receive direct inputs from the midbrain ^14,15^. Moreover, we modeled the afferent cortical projections to the STR as a unilateral connection from vmPFC to STR ^37^. Additionally, we excluded connections between regions lacking direct monosynaptic connections ^38^. We used fixed-effects Bayesian Method Selection (BMS) at the family level that showed that family [i], with a unilateral connection between vmPFC and STR represented the best explanation of the data with a total posterior probability of 0.99 as opposed to family [ii], [iii] and [iv] (posterior probability = 0.01, 0.0 and 0.0, Fig. 6a and 6b).

**Figure 6:**
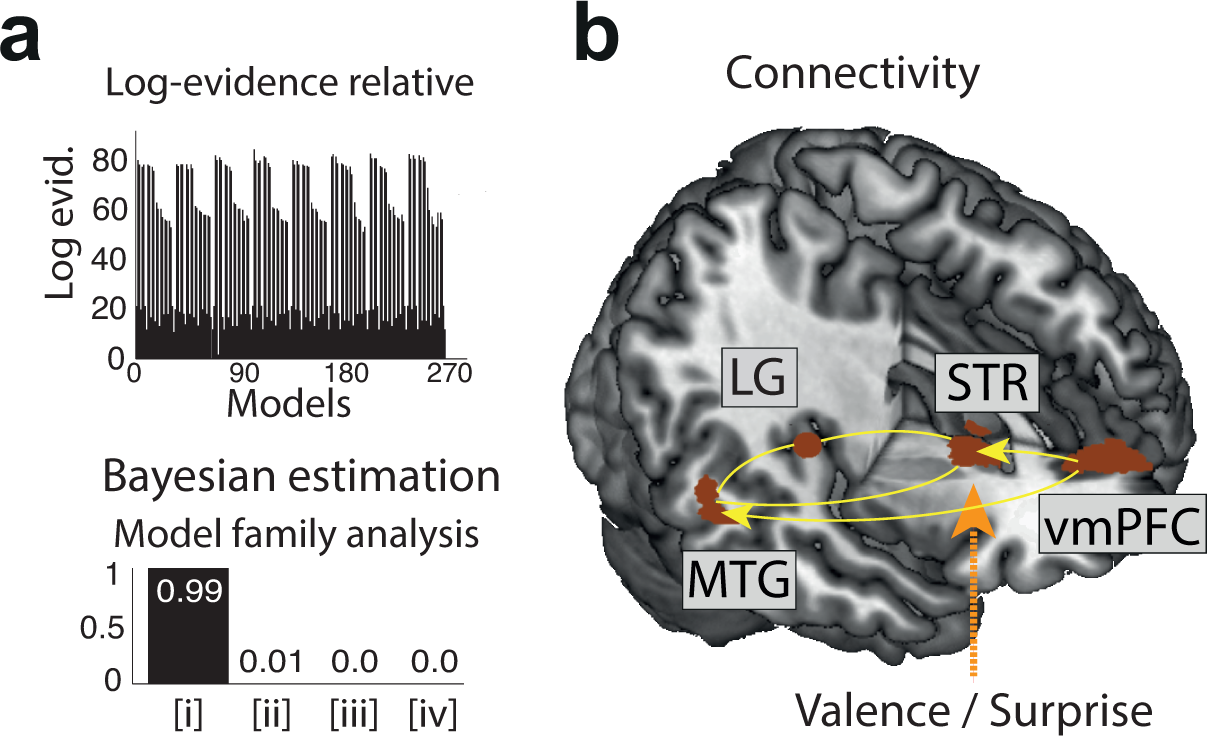
DCM results for the network encoding both outcome valence and surprise. (**a**) Bayesian estimation analyses revealed that the family of models with a unilateral connection between vmPFC and MTG (family [i]) was the best family (P > 0.99) as compared with three other possible families showing different effective connectivity between vmPFC and MTG. (**b**) The winning family of models was identified as the most likely of all the families of models representing the possible connections between the four areas found to encode both outcome valence and surprise. This family contains unilateral connections such that the vmPFC projects to both the MTG and STR with additional connections linking the STR to the MTG and LG with a double driving modulatory input (e.g. late outcome valence and surprise regressors) targeting the STR.

## Discussion

Here we provided a comprehensive spatiotemporal characterization of the neural correlates of outcome valence and surprise in the human brain by capitalizing on the high temporal and high spatial resolution of simultaneously acquired EEG and fMRI measurements. More specifically, by identifying temporally-specific EEG components of outcome valence and surprise and using the trial-by-trial amplitude fluctuations in these components as predictors in an fMRI analysis, we were able to demonstrate temporally overlapping but largely spatially separate representations for the two outcome dimensions. Overall, our work adds to the large body of related fMRI work ^7,19,39,40^ by offering a more complete spatiotemporal representation of the relevant networks.

As we demonstrated recently ^11^, electrophysiological variability in an outcome valence component was found to differentially suppress or activate regions of the human reward network in response to negative and positive outcomes respectively, consistent with a role in guiding approach and avoidance learning respectively ^11,34,41^. To examine the spatial extend of outcome surprise relative to this outcome valence representation, we used separate parametric fMRI predictors for unsigned RPE that were derived from a RL model and from the endogenous STV in a relevant EEG component. Our conventional model-based fMRI predictor revealed a distributed network of activations including the dlPFC, aINS and mPFC in line with previous reports in the literature ^12,22,23,31^. However, recent related findings claim that activity in regions other than those found in the conventional unsigned RPE network such as the STR or the vmPFC, might also encode outcome surprise information at time of outcome ^19,31,42,43^. Similarly, others have endorsed the notion of a ventral-dorsal gradient in the striatum encoding valence-surprise RPE information^23,44^.

Consistent with these recent claims, our EEG-informed outcome surprise regressor exposed additional unique areas, including the dorsal STR, the vmPFC and the lOFC among others. This finding endorses the hypothesis that trial-by-trial variability in our electrophysiologically-defined measure of outcome surprise (i.e. endogenous variability) may carry additional explanatory power compared to its behaviorally derived counterpart (i.e. external variability). It also highlights the utility of the simultaneous EEG-fMRI measurements in exposing latent brain states, thereby complementing more conventional model-based fMRI analyses.

Overall, these new results imply that surprise is encoded near simultaneously with outcome valence but in largely separate systems, reinforcing the notion that these outcome dimensions subserve related but separate functions. More specifically, our results are consistent with the idea that the late system encoding outcome valence determines the direction in which learning occurs (approach or avoidance learning), as it is up- and down-regulated following positive and negative outcomes respectively. In contrast, the system encoding outcome surprise captures the absolute discrepancy in stimulus-reward associations, thus being responsible for speeding up or slowing down the learning process ^31^. In fact, the outcome surprise network included many regions associated with the human attentional network, consistent with an increase in resource allocation for unexpected outcomes in order to facilitate learning^23,31,42^.

Despite the seemingly separate spatial representations associated with the outcome valence and surprise components we also found a spatial overlap in a smaller network comprising the STR, vmPFC, MTG and LG, with activity in these regions being predictive of stimulus value updating. Importantly, this overlap appeared exclusively in the conjunction of activations appearing in the EEG-derived measures of valence and surprise. While a large body of evidence have implicated the STR and vmPFC in reward learning ^30,45^ previous work in primates and humans offered contradicting evidence on whether these regions encode valence or unsigned RPE ^46,47^. Our results are intriguing as they reconcile previous reports and provide compelling evidence implicating the STR and vmPFC in processing both outcome valence and surprise.

In addition, our results provide further evidence that additional areas such as the MTG and LG are implicated in processing reward-based learning. This finding is consistent with some studies offering converging evidence that activity in MTG and LG signals behavioral switches in choice in non-primates ^48,49^ and in humans ^2,45^. However, another interpretation would be that activity in these visuo-mnemonic areas facilitate effective inference in changing environments ^10,50^ by modulating the stimulus-reward association established in memory ^45^. Here, our results confirm the role of these two regions in encoding stimulus value updating during learning and thus facilitating adaptive decisions.

Moreover, the fMRI signal in all four regions (STR, vmPFC, MTG and LG) exhibited a clear linear superposition profile of the two outcome-related signals. This profile is consistent with recent theories postulating a single population of neurons encoding an integrated representation of both signals in these regions ^51^. Alternatively, this finding could suggest the presence of separate but *intermixed* populations of outcome valence and surprise coding neurons in the network. Although speculative, this hypothesis would predict that the activity of different populations of neurons randomly distributed within an area is averaged out within fMRI voxels to give rise to the response profile we observed here. Recent evidence from electrophysiological investigations in animals supports this idea and demonstrates that valence and surprise signals are encoded by an equivalent amount of intermixed DA neurons with orthogonal coding patterns in the midbrain ^14,52-54^ and in the posterior parietal cortex, including the MTG ^14,15,22^. Supporting this idea, our connectivity analysis confirmed that the best neuronal model explaining our data required a double parallel driving input to the network (through the STR) encoding both outcome valence and surprise.

Similarly, this result is noteworthy because it provides the first evidence in humans, to our knowledge, that a composite signal – that does not rely entirely on a single RPE representation signal encoded by one family of dopamine (DA) neurons ^6,55-57^ – is necessary to drive learning. This interpretation is further supported by recent evidence failing to uncover a true RPE in the STR, especially when categorical outcome valence is accounted for ^58^, highlighting the collinearity confound between the two signals in conventional fMRI designs. Instead, our results are more consistent with another recent study reporting multiple outcome-related signals converging onto the STR ^59^.

In addition to receiving outcome valence and surprise signals from regions encoding primary reinforcement signals such as the midbrain and prefrontal cortex, our connectivity results suggest that the STR also interacts with visuo-mnemonic areas (such as the LG and MTG) in order to compute an integrative learning signal ^10,60^. Remarkably, the vmPFC also appears to transfer information to the MTG, directionally consistent with previously reported bold> effects of learning related signals in MTG ^61,62^ such that value information can be attributed to the visual aspect of the stimuli under consideration ^45,50^. Thus existing visual memories of the stimuli could be reactivated at time of outcome and updated with new rewarding information in order to help the decision maker select the optimal alternative in subsequent trials ^47,63^. Confirming this hypothesis, in a supplementary analysis, we found that the same four areas were parametrically modulated by the value of the chosen option in the decision phase, in line with previous reports ^41,64,65^

In summary, our data advance our understanding of the neurobiological mechanisms of learning associations between stimuli and outcomes and suggest complementary roles for outcome valence and surprise in decision making that can help constrain formal learning theories.

## Materials and Methods

### Participants

Twenty-four participants took part in the study, which was conducted in accordance with approved guidelines. Informed consent was obtained from all participants while all experimental protocols were approved by the School of Psychology Ethics Committee at the University of Nottingham. Four participants were excluded due to excessive head movements inside the scanner. The remaining twenty participants (12 females; average ± SD age, 21 years ± 2.6 years) were used in all subsequent analyses. All were right handed, had normal or corrected-to-normal vision and reported no previous history of neurological problems.

### Stimuli

Our task employed twelve abstract stimuli (examples given in Fig. 1a) and two feedback symbols (a tick and a cross for positive and negative feedback respectively). All stimuli (180×180 pixels) including those used for the feedback (125×125 pixels) and the fixation cross (30x30 pixels) were equated for luminance and contrast. The task was programmed with the Presentation software (Neurobehavioral Systems Inc., Albany, CA), presented using a computer running Windows Professional 7 (64bit, 3GB RAM, nVidia graphics card) and projected onto a screen placed 2.3 m from the participants (EPSON EMP-821 projector; refresh rate: 60Hz, resolution: 1280×1024pixels, projection screen size: 120x90cm’s). The stimuli and feedback symbols were subtended 4^°^ × 4^°^ and 3^°^ × 3^°^ of visual angle respectively.

### Probabilistic reversal learning task

The experiment consisted of two blocks of 170 trials each, separated by a break. At the beginning of each block, participants were shown three symbols (A, B and C) chosen randomly from the larger set of twelve stimuli. The chosen symbols were used throughout the block. In each trial, participants were told to identify the symbol with the highest reward probability among a pair of stimuli selected from the three symbols. Each rewarded trial earned them 1 point, while unrewarded trials earned them zero points. Participants knew that their performance would be monitored and transformed into monetary rewards at the end of the experiment (up to a maximum of £45), without being instructed on the exact mapping between earned points and their final payoff.

At any given point during the experiment, one of the three symbols was associated with a 70% chance of obtaining a reward (“high” reward probability symbol) compared to the remaining two symbols, each of which had a 30% chance of obtaining a reward (“low” probability symbols). Participants were not informed of the exact reward probabilities assigned to the symbols and they were told instead to learn to choose the symbol that was more likely to lead to a reward through trial and error (i.e. making use of the outcome of past decisions). To prevent participants from searching for non-existent patterns and to reduce cognitive load we presented the three possible pair combinations of the three symbols in a fixed order (i.e. AB, BC and CA) – though the presentation on the screen (left or right of the fixation cross) for the two symbols was randomized. Participants were explicitly informed about this manipulation.

Each trial began with the presentation of a fixation cross for a random delay (1-4 s; mean 2.5 s). To minimize large eye-movements, participants were instructed to focus on the central fixation. Two of the three symbols were then placed to either side of the fixation cross for 1.25 s. During this period, participants had to press a button on a fORP MRI compatible response box (Current Design Inc., Philadelphia, PA, USA) using either their right index or middle finger to select the right or left symbol on the screen, respectively. The fixation cross flickered for 100 ms after each button press to signal to the participants that their response was registered. Finally, the decision outcome was presented after a second random delay (1-4 s; mean 2.5 s). Rewarded or unrewarded feedback was given by placing a tick or a cross, respectively, in the center of the screen for 0.65 s. Trials, in which participants failed to respond within the 1.25 s of the stimulus presentation, were followed by a “Lost trial” message and were excluded from further analyses. To increase estimation efficiency in the fMRI analysis, the timing of the two delay periods was optimized using a genetic algorithm ^66,67^. Figure 1a summarizes the sequence of these events.

We defined a learning criterion for which participants were thought to have learned the task when they consistently selected the high probability symbol in 5 out of the last 6 trials. Every time this learning criterion was reached, we introduced a reversal by randomly re-assigning the “high” reward probability to a different symbol. This manipulation ensured that participants only experienced reversals after learning, as previously used in model-based fMRI studies ^25,68^. To make reversals less predictable, we included additional buffer trials after the learning criterion was reached. The number of buffer trials followed a Poisson process, such that there was a probability of 0.3 that a reversal occurred on any subsequent trial (with a minimum of 1 and a maximum of 8 trials). Finally, a key component of this paradigm was that each stimulus pair was chosen from a larger set of three symbols. This manipulation encouraged participants to engage in an exploration phase to identify the most rewarding symbol after reversals and it forced the participants to choose between the two least rewarding symbols even after they had learned the task (when the two were presented together). This manipulation ensured a more balanced number of positive and negative outcomes across all trials.

**Figure 1:**
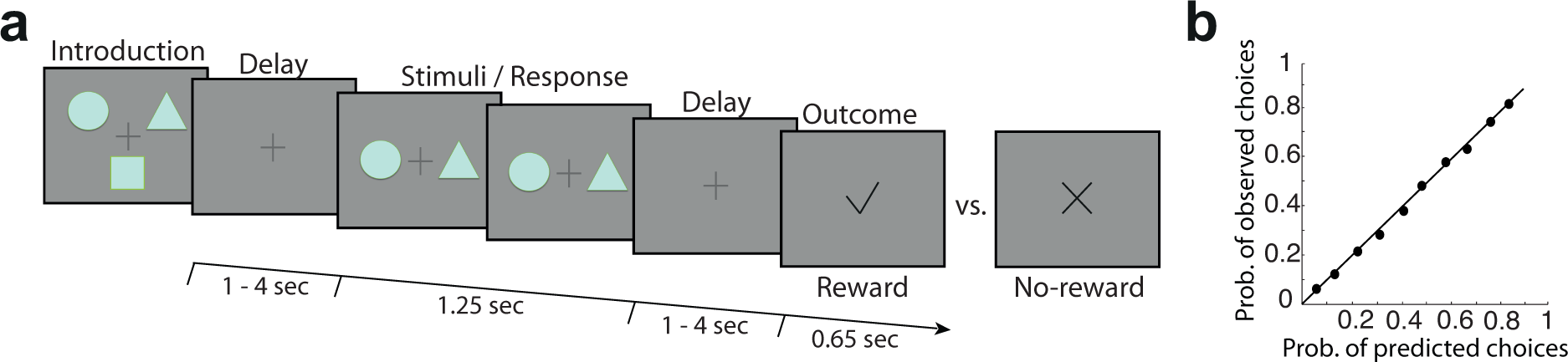
Schematic representation of the experimental task and modelling of behavioral responses. Each trial began with a random delay followed by the presentation of two abstract symbols (selected from a larger set of three symbols) for a period of 1.25 s. During this time, subjects pressed one of two buttons on a response device to indicate which of the two symbols (right or left) they believed was more likely to lead to a reward. The fixation cross flickered for 100 ms when a selection was made. Finally the decision outcome was revealed after a second random delay. A tick or a cross were used to inform the participants of a positive or a negative outcome, respectively.

### Modeling of behavioral data

We used a model-free reinforcement learning (RL) algorithm to estimate trial-by-trial RPEs using each participants’ behavioral choices ^1^. Specifically, the algorithm assigned each choice *i* (for example selecting the symbol A) an expected value *v*_A_(*i*) which was updated via a RPE, *δ*(*i*), as follows:

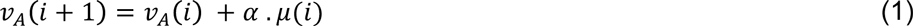

where α is a fixed learning rate that determines the influence of the RPE on the updating of the stimulus expected value. The RPE was given by the following equation: δ(*i*) = *r*(*t*) – *V*_A_(*i*), where *r*(*i*) represents the observed outcome (0 or 1). The expected values of both the unselected stimulus (e.g. B) and the stimulus not shown on trial *i* (e.g. C) were not updated.

It is worth noting that subject-wise differences in overall task volatility (contingent upon the number of reversals attained during the task) were captured by different subject-wise estimates of the learning rate (for example, subjects who experienced a more stable environment – that is fewer reversals – had lower learning rate estimates). For comparison, to capture both subject- and trial-wise fluctuations in task volatility we also used a RL model incorporating a dynamic learning rate (DYNA) that enables a trial-by-trial scaling of the choice expected value updating (Krugel et al., 2010). In this model the learning rate (α) on each trial *i* is modulated by the slope of the smoothed unsigned RPE (*m*) according to the following update equations:

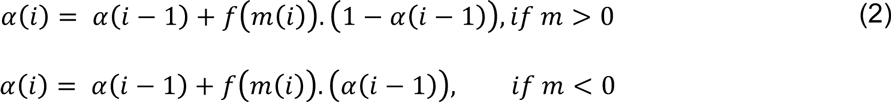

where *f*(*m*(*i*) is a double sigmoid function that transforms the slope *m* into the [0,1] interval and scales the trial-by-trial updating of the dynamic learning rate. Crucially, this transformation function is itself parameterized by a free parameter γ. High values of γ render the updating of the dynamic learning rate negligible so that in essence the learning rate becomes fixed. On inspection of the subject-wise fits of the dynamic learning rate model we found high values of the parameter γ across participants and unvarying trial-by-trial estimates of the dynamic learning rate (across subjects SD = 0.01).

Whilst model-free RL approaches (such as the ones presented above) rest on the assumption that subjects make choices contingent upon the cached stimulus-expected value associations that have been acquired through prior experience, model-based RL approaches allow for representations of stimulus-outcome contingencies to bear on the decision process. To rule out that our subjects were merely inferring stimulus-outcome contingencies instead, we adapted the model presented in ^33^ to our one stage task environment. Briefly, in our model-based RL model stimulus-outcome contingencies were updated as follows: *SO*(*i* + 1,*s*,*r*) = *SO*(*i*,*s*,*r*) + αδ(*i*) where SO represents a stimulus-outcome contingency matrix, *s* indicates the chosen stimulus, *r* specifies the type of binary outcome (1 for positive outcome and 0 for negative outcome), α is a fixed learning rate and δ is a stimulus-outcome prediction error computed as δ(*i*) = 1 − *SO*(*i*,*s*,*r*). The stimulus-outcome contingencies of the two stimuli that were not chosen and not shown on trial *i* were not updated.

As in the model-free variants of our RL model only the expected value of the chosen stimulus (e.g. A) was updated as *v*_A_(*i* + 1) = SO(*i* + 1,A,1). In all models we used a softmax decision function in which, on each trial *i*, a stimulus choice probability (e.g. A) was given by:

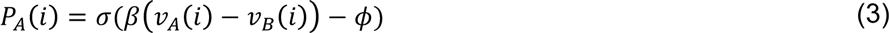

where σ(*z*) = 1/(1 + *e*^−*z*^) is the logistic function, *ϕ* denotes the indecision point (when choosing each of the alternatives is equiprobable) and *β* represents the degree of choice stochasticity (i.e., the exploration/exploitation parameter). Whilst choice probability of the unchosen stimulus (e.g B) was updated as follows: *P*_B_(*i*) = 1 − *P*_A_(*i*), choice probability of the stimulus not shown on trial *i* (e.g. C) was not updated.

### Model fitting and model comparison

For each subject *j* we estimated the set of model parameters *θ*_*J*_ *using a maximum likelihood estimation* fitting procedure: 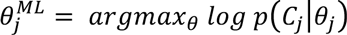 where *log p*(*C*_*j*_|*θ*_*j*_) is the choice log-likelihood given the model parameters *θ*_*j*_. To preserve the parameters’ natural bounds, log (*β*,*γ*) and logit (*α*) transforms of the parameters were implemented.

To determine the best fitting model we performed classical model comparison. Specifically, for each model we first estimated the subject-wise Bayesian Information Criterion (BIC) as follows: *BIC* = −2*log p*(*C*_j_|*θ_j_*) *+ dlogn*. Here, the goodness of fit of a model (−*log p*(*C*_*j*_|*θ*_*j*_)) is penalised by the complexity term (*dlogn*) where the number of free parameters in the model *d* is scaled by the number of data points *n* (i.e. trials). We then computed the sum of subject-wise BIC for each model and compared the model-wise BIC estimates (lower estimates indicating better fit). We found the BIC for the model-free RL with a fixed learning rate to be the lowest (*BIC*_*MF*_ = 413.26; *BIC*_*MB*_ = 413.61; *BIC*_*DYNA*_ = 529.17).

To visualize the winning model goodness of fit we divided the subject-wise predicted choice probabilities into five groups depending on the distribution quintiles and for each group computed the subject-wise average predicted choice probability. Using the sorting index of the predicted choice probabilities we then retrieved the subject- and group-wise average observed choice probabilities and computed Pearson’s correlation coefficient (rho = 0.97, Fig. 1b).

Finally, we used the RPE estimates from the model-free RL with a fixed learning rate to provisionally separate trials into those with low-vs-high RPEs surprise to run a binary decoder on the EEG data (see below). Our main goal was to then exploit the single-trial variability in the electrophysiologically-derived (i.e. endogenous) and temporally-specific representations of outcome valence and surprise to build fMRI predictors in order to identify the networks associated with these representations in the human brain.

### Electrophysiological recordings

EEG recordings were performed simultaneously with the fMRI acquisition. EEG data were acquired using Brain Vision Recorder (BVR; Version 1.10, Brain Products, Germany) and MR-compatible amplifiers (BrainAmps MR-Plus, Brain Products, Germany). The sampling rate was set at 5 kHz with a hardware bandpass filter of 0.016 to 250 Hz. The data were collected with an electrode cap consisting of 64 Ag/AgCl scalp electrodes (BrainCap MR; Brain Products, Germany) placed according to the international 10–20 system. Reference and ground electrodes were embedded within the EEG cap and positioned along the midline (reference: placed between electrodes Fpz and Fz, ground electrode: placed between electrode Pz and Oz). 10 kΩ surface-mount resistors were placed in line with each electrode for added subject safety, while all leads were twisted for their entire length and bundled together to minimize inductive pick-up. All input impedances were kept below 20 kΩ. Hardware-based EEG/MR clock synchronization was achieved using the SyncBox device (Syncbox, Brain Products, Germany) and MR-scanner triggers were collected to enable offline removal of MR gradient artifacts (GA). Scanner trigger pulses were stretched to 50µs using an in-house pulse extender to facilitate accurate capture. These pulses along with all experimental event triggers were recorded by the BVR software to ensure synchronization with the EEG data.

To minimize the influence of the GA at source, electrodes Fp1 and Fp2 were positioned at the scanner’s z = 0 position (i.e. the scanner’s isocenter), corresponding to a 4 cm shift of the head in the foot-head direction ^69^. This was achieved by aligning electrodes Fp1 and Fp2 with the laser beam used to position the subject inside the scanner. We used a 32-channel SENSE head coil with an access port that allowed all EEG cables to run along a straight path out of the MR, thereby ensuring no wire loops and minimizing the risk of RF heating the EEG cap. Finally we used a cantilevered beam to isolate the cables from scanner vibrations, thereby reducing induced artifacts as much as possible ^70^.

#### EEG pre-processing

We designed offline Matlab (Mathworks, Natick, MA) routines to perform basic processing of the EEG signals and remove GA and ballistocardiogram (BCG) artifacts (appearing due to magnetic induction on the EEG leads). The GA is inherently periodic and we therefore created an artifact template by averaging the EEG signal over 80 functional volumes and subsequently subtracting it from the EEG data centered on the middle volume. This procedure was repeated for each volume and each EEG channel separately. To remove any residual spike artifacts, we applied a 10 ms median filter. We then applied (non-causally to avoid distortions caused by phase delays) the following filters: a 0.5 Hz high-pass filter to remove DC drifts, 50 Hz and 100 Hz notch filters to remove electrical line noise, and 100 Hz low pass filter to remove high frequency signals not associated with neurophysiological processes of interest.

Finally, we treated the BCG artifacts, which often share frequency content with the EEG. In order to avoid loss of signal power that might otherwise be informative we adopted a conservative and previously validated approach ^11,28^. Specifically, we only removed few subject-specific BCG components using principal component analysis and relied on our single-trial discriminant analysis (see below) to identify components that were likely to be orthogonal to the BCG (this is ultimately achieved due to the multivariate nature of our discriminant analysis). To extract the BCG principal components, we first low-pass filtered the EEG data at 4 Hz (e.g. the frequency range where BCG artifacts are observed), and then estimated subject-specific principal components. The average number of components across subjects was 2.3. The sensor weightings corresponding to the BCG components were then projected onto the broadband data and subtracted out.

#### Eye-movement artifact removal

Before performing the main task, our participants were asked to complete an eye movement calibration task including blinking repeatedly when a fixation cross appeared in the center of the projection screen and making several horizontal and vertical saccades depending on the position of a fixation cross, subtending 0.6^°^ × 0.6^°^ of visual angle. Using principal component analysis we determined linear EEG sensor weightings corresponding to eye blinks and saccades such that these components were projected onto the broadband data from the main task and subtracted out ^28,71^.

#### EEG linear discriminant analysis

We used single-trial multivariate discriminant analysis of the EEG ^11,26,72,73^ to perform binary discriminations along the outcome valence and surprise dimensions. Specifically, for each participant, we estimated linear weighting of the EEG electrode signals (i.e. spatial filters) that maximally discriminated between 1) positive vs. negative feedback trials (valence dimension) and 2) high vs. low outcome surprise trials (surprise dimension) (Eq. 5). To identify temporally distinct neuronal components associated with each dimension we employed this procedure over several temporally distinct training windows. Applying the estimated spatial filters to single-trial data produced a measurement of the resultant discriminating component amplitudes. These values represent the distance of individual trials from the discriminating hyperplane and can be thought of as a surrogate for the neuronal response variability associated with each discriminating condition, with activity common to both conditions removed (Fig. 2a) ^72^.

Here, we defined discriminator training windows of interest with width 60 ms and center times τ, ranging from −100to 600 ms relative to outcome onset (in 10 ms increments) and used a regularized Fisher discriminant to estimate a spatial weighting vector ***w***(τ), which maximally discriminates between sensor array signals ***x***(*t*), for two conditions (i.e. positive vs negative outcomes and high vs low surprise RPE trials):

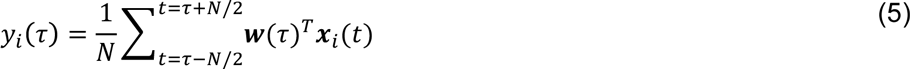

where ***x***(*t*) is an D × T matrix (D sensors and T time samples) and *y*_*i*_(τ) represents the resulting single-trial discriminator amplitudes. In separating the relevant groups of trials for outcome valence and surprise the discriminator was designed to map positive outcome and high surprise RPE trials to positive amplitudes and negative outcome and low surprise RPE trials to negative amplitudes.

Our aim was to capitalize on the endogenous variability in these single-trial discriminator amplitudes (*y*_*i*_(τ)) to build EEG-informed BOLD predictors (Fig. 2b) for analyzing the simultaneously acquired fMRI data (see below). Our working hypothesis is that single-trial variability (STV) in our electrophysiologically-derived measures of outcome valence and surprise can enable otherwise “static” fMRI activations (resulting from temporal averaging and the slow dynamics of conventional fMRI) to be absorbed by temporally specific components, thereby offering a more complete spatiotemporal picture of the underlying networks. Note that while deeper and subcortical structures contribute less to the EEG signal, our approach can still expose these regions as long as they covary with the cortical sources of the EEG STV.

We estimated the spatial vectors ***w***(τ) in Equation 5 for each time window τ as follows: ***w*** = *S*_*c*_(*m*_*2*_ – *m*_*1*_) where *m*_*i*_ is the estimated mean for condition *i* and ***S***_*c*_ = 1/2(***S***_1_ + ***S***_2_) is the estimated common covariance matrix (i.e. the average of the empirical covariance matrices for each condition, 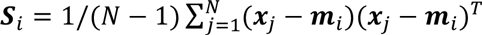 with *N* = number of trials). To treat potential estimation errors we replaced the condition-wise covariance matrices with regularized versions: 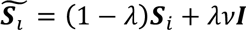 with λ ϵ [0,1] being the regularization term and *v* the average eigenvalue of the original *S*_*i*_ (i.e. trace(*S*_*i*_)/D). Note that λ = 0 yields unregularized estimation and λ = 1 assumes spherical covariance matrices. We optimized λ for each subject separately during the entire period following the outcome, using a leave-one-out trial cross validation (λ’s, mean ± se: 0.028 ± 0.05).

To quantify the discriminator’s performance at each time window, we calculated the area under a receiver operating characteristic curve, also known as *Az* value, using a leave-one-out trial cross validation ^74^. Next, we assessed the significance of the discrimination performance using a permutation test by performing the leave-one-out trial procedure after randomizing the labels associated with each trial. To produce a probability distribution for *Az*, and estimate the Az value leading to a significance level of P < 0.01, we repeated this randomization procedure 1000 times. Note that our classification results were virtually identical when using a logistic regression approach to train the classifier ^27^.

Given the linearity of our approach, we also computed scalp topographies for each discriminating component of interest resulting from Equation 5 by estimating a forward model as:

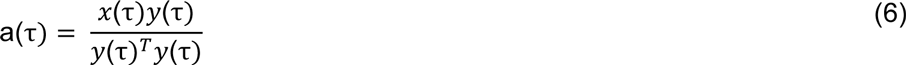

where *y*_*i*_(*λ*) is now shown as a vector ***y***(λ), where each row being a trial, and ***x***_*i*_(*t*) is organized as a matrix, ***x***(λ), with rows as channels and columns as trials, all for time window λ. These forward models can be viewed as scalp plots and interpreted as the coupling between the discriminating components and the observed EEG ^11,27,28,73^.

#### MRI data acquisition

MRI data were acquired on a 3 Tesla Philips Achieva MRI scanner (Philips, Netherlands). Our functional MRI data were collected using a 32-channel SENSE head coil (SENSE factor = 2.3) measuring 40 slices of 80×80 voxels (3 mm isotropic), a field of view (FOV) of 204 mm, flip angle (FA) of 80^°^, repetition time (TR) of 2.5 s and echo time (TE) of 40 ms. Slices were acquired in interleaved fashion. In total, two runs of 468 functional volumes, corresponding to the blocks of trials in the main task, were acquired. To correct for B0 inhomogeneities in the fMRI data, a B0 map was also acquired using a multishot gradient echo sequence (32 slices of 80×80 voxels – 3 mm isotropic, FOV: 204 mm, FA: 90^°^, TR: 383 ms, TE: 2.3ms, delta TE: 5ms). Anatomical images were collected using a MPRAGE T1-weighted sequence (160 slices of 256×256 voxels – 1mm isotropic, FOV: 256 mm, TR: 8.2 ms, TE: 3.7 ms).

#### fMRI data preprocessing

The first five volumes from each experimental run were removed to ensure that steady-state imaging has been achieved. We used the remaining 463 volumes for all subsequent statistical analyses. fMRI data preprocessing included motion correction, slice-timing correction, high-pass filtering (>100 s) and spatial smoothing (with a Gaussian kernel of 8 mm full-width at half maximum). These steps were achieved using the FMRIB’s Software Library (Functional MRI of the Brain, Oxford, UK). Next, we applied B0 unwarping onto the fMRI images to correct for signal loss and geometric distortions due to B0 field inhomogeneities. Finally, the registration of fMRI data to standard space (Montreal Neurological Institute, MNI) was performed using FMRIB’s Non-linear Image Registration Tool using a 10 mm warp resolution. This procedure included an initial linear transformation of the fMRI images into an individual’s high-resolution space (using six degrees of freedom) prior to applying the non-linear transformation to standard space.

#### fMRI analysis

We employed a multilevel approach within the general linear model (GLM) framework to perform whole-brain statistical analyses of functional data as implemented in FSL ^75^:

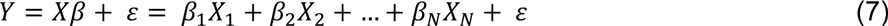

where *Y* is a *T* × 1 (T time samples) column vector containing the times series data for a given voxel, and *X* is a T × *N* (*N* regressors) design matrix with columns representing each of the psychological regressors convolved with a canonical hemodynamic response function (double-*γ* function). β is a *N* × 1 column vector of regression coefficients and ε a *T* × 1 column vector of residual error terms. Using this model we initially performed a first-level fixed effects analysis to process each participant’s individual experimental runs, which were then combined in a second-level fixed effects analysis. Finally, we combined results across participants in a third-level, mixed-effects model (FLAME 1), treating subjects as a random effect. Time series statistical analysis was carried out using FMRIB’s improved linear model with local autocorrelation correction. Applying this framework, we performed the GLMs highlighted below.

#### GLM1 – EEG-informed fMRI analysis of outcome phase

Our main fMRI analysis was designed to reveal the brain networks underlying the two main outcome dimensions – valence and surprise – by capitalizing on the endogenous STV in temporally-specific EEG components associated with these dimensions. For the valence dimension, we aimed at exposing brain areas in which the BOLD signal varied with the EEG STV along a positive versus negative axis, while for the surprise dimension the BOLD signal varied with the EEG STV along a high versus low surprise RPE axis, regardless of valence. Specifically, locked to the time of outcome (i.e. when the tick/cross appeared) we included five boxcar regressor with a duration of 100 ms for each regressor event: 1) an unmodulated regressor (UM; all event amplitudes set to one), 2) a fully parametric regressor whose event amplitudes were modulated by the EEG STV associated with the outcome valence component of interest (*EEG*_*LateVa1*_), 3) a fully parametric regressor whose event amplitudes were modulated by the outcome surprise estimates from the RL model (*MODEL*_*Mag*_), and 4) a fully parametric regressor whose event amplitudes were modulated by the EEG STV associated with a RPE component of interest (*EEG*_*LateMag*_). Finally, we included a fully parametric regressor whose event amplitudes were modulated by the EEG STV associated with an earlier outcome valence component (to account for fast alertness response to negative outcomes as per ^11^, *EEG*_*EarlyVal*_), an unmodulated regressor for all lost trials at time of outcome (*LOST*), an unmodulated regressor of no interest at the time of stimulus presentation (i.e. decision phase, *DEC*), and six nuisance regressors, one for each of the motion parameters (three rotations and three translations), such as:

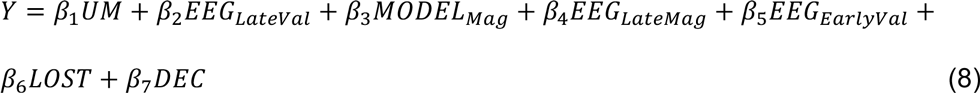

Note that the amplitudes of the two valence regressors (*EEG*_*ValLate*_ and *EEG*_*VaiEarly*_) were largely uncorrelated ^11^.

To examine the extent to which neural representations of outcome valence and surprise overlap spatially across participants, we performed a conjunction analysis on the results arising from the EEG-informed regressors for the late valence and surprise components. Specifically, we examined which brain areas were jointly activated by creating an intersection of statistical maps relating to the two components (as implemented in *easythresh_conj* within FSL ^76^). The conjunction analysis was performed at the whole-brain level and the resulting statistical image was thresholded at Z-score > 2.57 with a cluster probability threshold of P = 0.05.

#### GLM2 – Region of interest analysis

To visualize the overall response profile of regions of interest (ROIs) (e.g. those representing both valence and surprise from GLM1) we ran another model with trials binned into six outcome-locked regressors, separated into positive and negative outcomes as well as according to three surprise groups for each outcome type (low, medium, high) using the EEG STV in our outcome surprise component (0-33%, 33-66%, and 66-100% percentiles of EEG amplitudes). The main motivation for this analysis was to visualize the response profile within these areas with respect to the strength of the endogenous variability carried by the EEG STV (rather than the variability from unsigned RPE amplitudes from the RL model). In addition we included the same regressors of no interest we used in GLM1 above.

From each of the six regressors we extracted beta coefficients from the following ROIs: ventromedial prefrontal cortex (vmPFC), striatum (STR), middle temporal gyrus (MTG) and lingual gyrus (LG). More specifically, we first extracted ROIs masks at the group level from GLM1 (i.e. in standard space), by applying the cluster correction procedure described above. We subsequently back-projected these ROIs from standard space into each individual’s EPI (functional) space by applying the inverse transformations as estimated during registration (see *fMRI data preprocessing section*). Each ROI was then checked against the relevant (regressor-specific) statistical maps in the individual brains [at a slightly more lenient threshold of P < 0.01 uncorrected, cluster size > 10 voxels (90 mm^3^)] to ensure that the inverse-transformation was performed properly. Finally, for each of the six regressors we computed average beta coefficients from all voxels in the back-projected clusters and across participants to visualize the overall response profile of the ROIs as a function of both outcome valence and surprise.

#### Relationship between percent signal change and value updating

To establish a more concrete link between the brain regions encoding both outcome-related signals, we ran an additional regression analysis. Specifically, we directly tested the relationship between trial-related percent signal change (PSC) in those areas and stimulus value updating, the key underlying component of learning, defined as the difference in successive prediction values, *v*^*k*^ (*i*+1) − *v*^*k*^(*i*), computed using estimates of our RL model, with *k* indexing a given choice and with i indexing trial. PSC was estimated on outcome-locked data by defining a temporal window extending 5 s (2 volumes) before the outcome and extending to 10 s (4 volumes) after the outcome onset. We then estimated the PSC traces for all outcomes as follows:

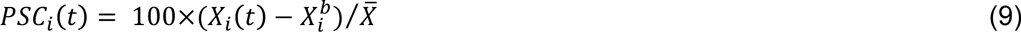

where *X*(*t*) is the mean outcome-locked data at time point *t, X*^b^ is the mean baseline signal defined as the average signal during the 5 s preceding each outcome and extending two volumes after onset, and 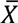 the mean outcome-locked volume signal across all data points in a given run. Finally, we employed the PSC as our predictor of value update in a linear regression such as:

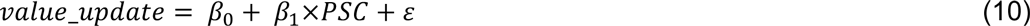

Then, to establish a significant trial-by-trial association between PSC in our ROIs and value update, we tested whether the regression coefficients resulting from all subjects (*β*_1_’s) come from a distribution with mean greater than zero (using a one-tailed *t*-test).

#### Resampling procedure for fMRI thresholding

To properly correct the statistical maps for multiple comparisons we employed a resampling procedure that considers the a priori statistics of the single-trial variability in all of our parametric predictors in a way that trades off cluster size and maximum voxel Z-score ^11^. Specifically, we preserved the overall distributions of the EEG discriminating components (*y*_*i*_(τ) for the outcome valence and surprise components) as well as the single-trial variability of the unsigned RPE predictor while removing the specific single-trial correlations in subject-specific experimental runs. Thus, for each resampled iteration and each predictor, all trials were built upon the original regressor amplitude distribution; however the specific values were mixed across trials and runs. In other words, the same regressor amplitudes were used in each permutation, however, the sequence of regressor events was randomized.

We repeated this resampling procedure 100 times for each participant, including a full three-level analysis (run, subject, and group). The design matrix included the same regressors of non-interest used in all our GLM analyses. Consequently, this process enabled us to construct the null hypothesis H0, and establish a joint threshold on cluster size and Z-score based on the cluster outputs from the randomized predictors. We then extracted cluster sizes from all activations with |Z|-score > 2.57 for both positive and negative correlations with the permuted predictors. Finally, we produced a distribution for the cluster sizes for the permuted data, and estimated the largest cluster size leading to a significance level of P < 0.05. This procedure resulted in a corrected threshold for our statistical maps, which we then applied to the activations observed in the original data. This cluster-threshold was set to > 76 voxels at |Z| = 2.57.

#### Exploratory causal analysis

To explore possible spatiotemporal models of outcome valence and surprise processing, we applied dynamic causal modelling (DCM) on our data using the DCM10 toolbox for SPM12 ^77^. Our goal was to explore potential causal relationships between the regions identified in the previous conjunction analysis (GLM1). The DCM analysis was carried out in several steps. We first entered the full GLM1 into SPM12 to generate a design matrix and extracted activation times courses for each region of interest from every participant. Secondly, we constructed a realistic model space covering an extensive (although non-exhaustive) series of causal models including possible combination of 1) the intrinsic connectivity between the nodes of interest during the course of the experiment and 2) the influence of the driving input, as per GLM1: the EEG STV representing outcome valence and surprise signals. DCMs are defined in terms of fixed (endogenous) connections between brains areas and input-specific changes in the strength of these connections (i.e. modulatory effects). In the present analysis, we focused on fixed connections between brain areas as we were interested in the potential causal relationships between the vmPFC, the STR, MTG and LG. On the basis of the principles governing RL models (i.e. *simultaneous* representation of outcome valence and surprise driving learning) and previous neurophysiological reports, the driving inputs representing both valence and surprise were added to the STR, which is known to receive direct inputs from the midbrain ^14,15^. Thus, in all models, the EEG STV related to outcome valence and surprise were fixed as a double driving input to the STR node. Moreover, we modeled the afferent cortical projections to the STR as a unilateral connection from vmPFC to STR ^37^. Additionally, we excluded connections between regions that have been shown to lack direct monosynaptic connections ^38^ such as the LG and the vmPFC.

We used a fixed-effects approach to Bayesian model selection to determine which family of models best explained our observed responses in vmPFC, STR, MTG and LG. This analysis assumes that all subjects engaged the same neuronal structure for processing outcome valence and surprise ^78^. Thus, we used family-level inference ^79^ and partitioned our model space including 256 models into four equally large subsets of models (e.g. families of 64 models) that differed in their endogenous connections such that family [i] represented all models in which the vmPFC had a unilateral connection with MTG such that vmPFC -> MTG, family [ii] represented all models in which the vmPFC had a unilateral connection with MTG such that MTG -> vmPFC, family [iii] represented all models in which the vmPFC had a bilateral connection with MTG (vmPFC <-> MTG) and family [iv] represented all models in which the vmPFC had no connection with MTG.

